# Variation in mate competition favors phenotypic plasticity in male coloration of an African cichlid

**DOI:** 10.1101/2022.11.15.516656

**Authors:** Robert J. Fialkowski, Tyler R. Funnell, Taylor J. Piefke, Shana E. Border, Phil M. Aufdemberge, Hailey A. Hartman, Peter D. Dijkstra

## Abstract

Sexual selection is thought to be a potent evolutionary force giving rise to diversity in sexual traits that enhance mating success, such as ornament, sexual display, and weapons. The expression of sexual traits is often influenced by environmental conditions, suggesting that phenotypic plasticity may precede and facilitate evolutionary divergence in sexual traits by sexual selection. However, the mechanisms that promote plasticity in sexual traits remain poorly understood. Using the cichlid fish *Astatotilapia burtoni*, we show that sexual selection may promote plasticity in sexual traits. In this species, males change between yellow and blue color and exhibit intense male contest competition over breeding territories to attract females. We found that experimentally increased competition over territories led to a higher proportion of males expressing the yellow phenotype. Furthermore, we found that expression of yellow coloration is beneficial to males in territorial disputes. Specifically, yellow males won more staged dyadic contests than blue males, and yellow males had a lower level of oxidative stress than blue males, especially when engaging in more territorial display. However, females spawned more with blue males than with yellow males, suggesting that expression of blue coloration is sexually more attractive. The ability to adjust color phenotype according to the local competitive environment could therefore promote the persistence of plastic changes in coloration. Our findings challenge the view that phenotypic plasticity sets the stage for the evolution of genetically fixed changes via sexual selection, and instead suggest that sexual selection may favor plasticity in sexual traits.

## Introduction

Competition for mates can drive diversification in sexual traits that enhance mating success, such as complex mating songs, bright color patterns, or weaponry (1, 2). The expression of these sexual traits is often dependent on environmental variation, which can give rise to phenotypic plasticity (3–5). Phenotypic plasticity is thought to play an important role in adaptive evolution (6–10), and recently, the possibility that phenotypic plasticity and sexual selection jointly drive diversification in sexual traits has received considerable attention (11–15). In animal taxa exhibiting extreme diversity in sexual traits, genetic differences in sexual traits found between species are often also found within species as environmentally induced phenotypic variants (8, 15). These findings suggest that phenotypic plasticity may be a steppingstone in adaptive divergence by sexual selection under certain circumstances. It is generally thought that ecological or abiotic factors, such as the signaling environment, temperature, or predation risk, drive phenotypic plasticity in sexual traits (16–18). Divergent sexual selection then refines the alternative sexual phenotypes, ultimately leading to genetic fixation of sexual traits. In this scenario, phenotypic plasticity is reduced due to strong sexual selection and the cost or constraints of plasticity (19, 20). However, the possibility that sexual selection promotes rather than reduces plasticity in sexual traits is rarely considered (11, 21, 22), and could have important consequences for our understanding of the interaction between phenotypic plasticity and sexual selection in adaptive evolution.

It is well known that changes in the social environment can impact sexual trait expression and alter the outcome of sexual selection (3, 23). Specifically, the intensity of male-male competition can vary widely at relatively short time scales due to environmental and demographic factors which can alter population density (24), operational sex ratio (25, 26), or availability of breeding territories (27). Under alternating low- and high-competition environments, selection could favor individuals that can reversibly adjust one’s phenotype according to the prevailing competitive environment, as has been suggested for the evolution of conditional strategies in relation to alternative mating tactics (28). Specifically, if alternative male phenotypes experience increased success in either outcompeting conspecific rivals or attracting mates, males should express the more competitive phenotype under periods of more intense male contest competition and the more sexually attractive phenotype when competition is less intense. This would in turn favor reversible phenotypic plasticity because it allows males to adjust their phenotype to the level of competition in their current environment.

Here we tested whether sexual selection favors phenotypic plasticity in a species that shows extreme plasticity in body coloration. The African cichlid fish *Astatotilapia burtoni* lives in shallow pools in East Africa and many populations consist of blue and yellow males (12, 29, 30), with males alternating repeatedly between the yellow and blue color phenotype^9^. As in other haplochromine cichlid species, male-male competition is intense in *A. burtoni* with males competing aggressively for territories, which are a prerequisite for reproductive success (27). Additionally, previous studies have indicated both color phenotypes utilize the same mating tactic (both compete aggressively for mating territories (31)).

Pigmentary differences between yellow males and blue males and color change involve both rapid physiological adjustments resulting from yellow pigment granule movement, and slower morphological changes in chromatophore density (32). Populations of *A. burtoni* with plastic yellow and blue coloration are common both in the wild and in the lab, but the factors that maintain plasticity between these two discrete color types remain elusive.

## Results

To test whether the intensity of male-male competition influences the expression of yellow and blue color phenotype, we manipulated the social environment by altering both the physical location and number of potential mating territories in replicate experimental communities during a 22-week period. This habitat manipulation mimics natural conditions where strong winds or hippopotami (*Hippopotamus amphibius*) can alter territorial structures in shallow waters (27). The habitat manipulation resulted in more social instability as evidenced by frequent changes in social status and heightened competition compared to control communities (**Extended Data Fig. 1**). Color change was common, with 84.7% of males changing color phenotype at least once with a median frequency of 3 color shifts during the 22-week period (**Fig. 1a, b**). The total number of color shifts was not influenced by instability treatment (GLMM, 0.073 ± 0.15, z = 0.504, *P* = 0.615). Next, we examined how social instability influences color expression. The proportion of yellow males increased over the 22-week experimental period at a higher rate in the instability treatment compared to the control (GLMM, effect of time x stability treatment: 0.0101 ± 0.0013, z = 7.610, *P* < 0.00001, **Fig. 2a**). Additionally, males that experienced more social status shifts were more likely to be yellow at the end of the experiment in both treatments (GLMM, effect of final color phenotype: 0.621 ± 0.167, z= 3.727, *P* = 0.000294, **Fig. 2b**). These results indicate that social instability promotes the expression of the yellow phenotype.

**Fig. 1.**
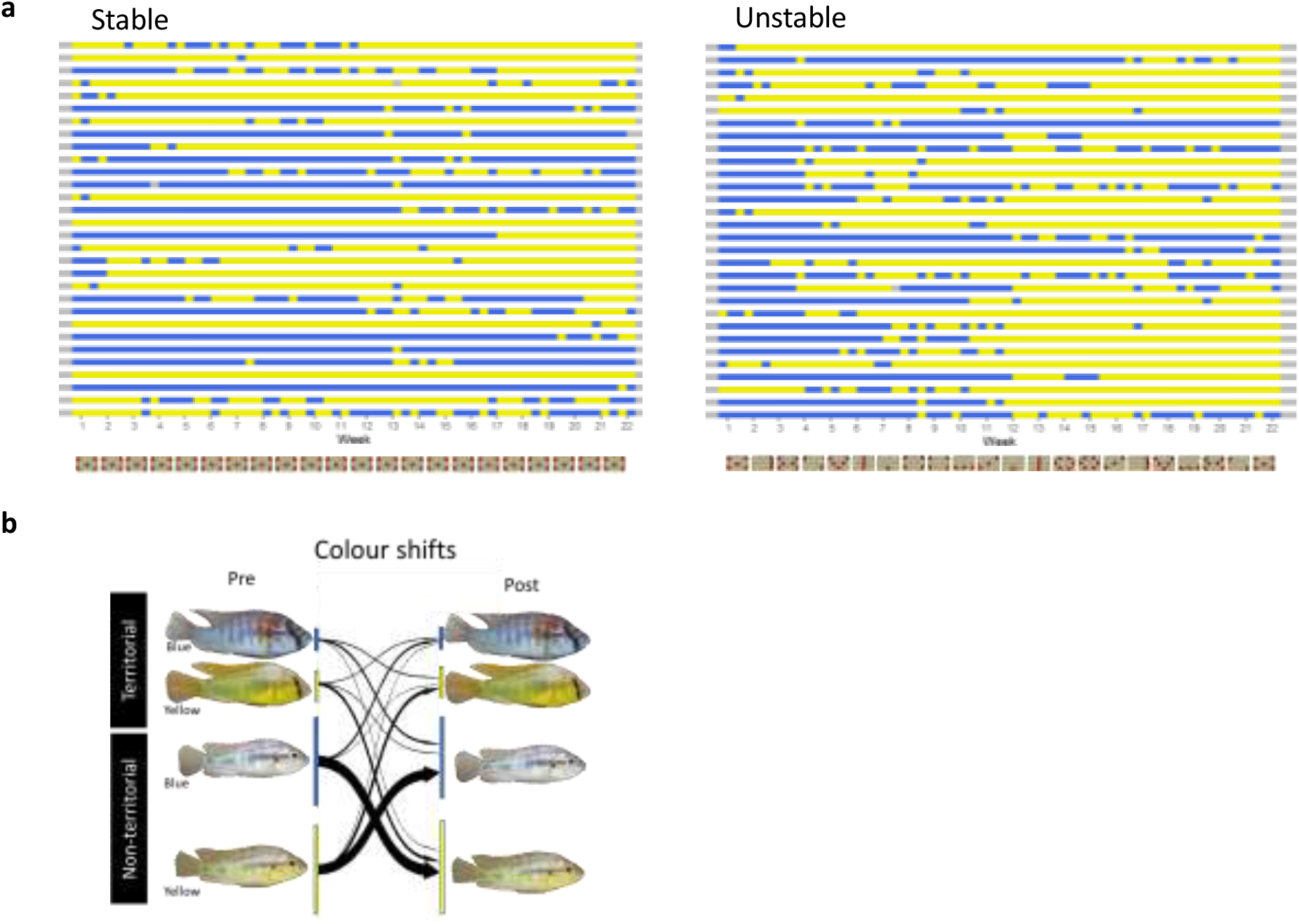
Color phenotype changes in stable and unstable communities. **a**, Graphical representation of color phenotype expression during the 22-week experiment in a subset of males from stable (left) and unstable communities (right). Each row represents an individual male. Shown at the bottom is a schematic representation of the arrangement of terracotta pot shards (top view) from week 1 to week 22. **b**, Color phenotype shifts in territorial and nonterritorial males during the experiment. A color phenotype shift is defined as a male expressing a different color phenotype between two consecutive observations (‘pre’ and ‘post’). The arrow thickness is proportional to the percentage of changes that occur during the entire duration of the experiment in both treatments combined (n = 144 males).

**Fig. 2.**
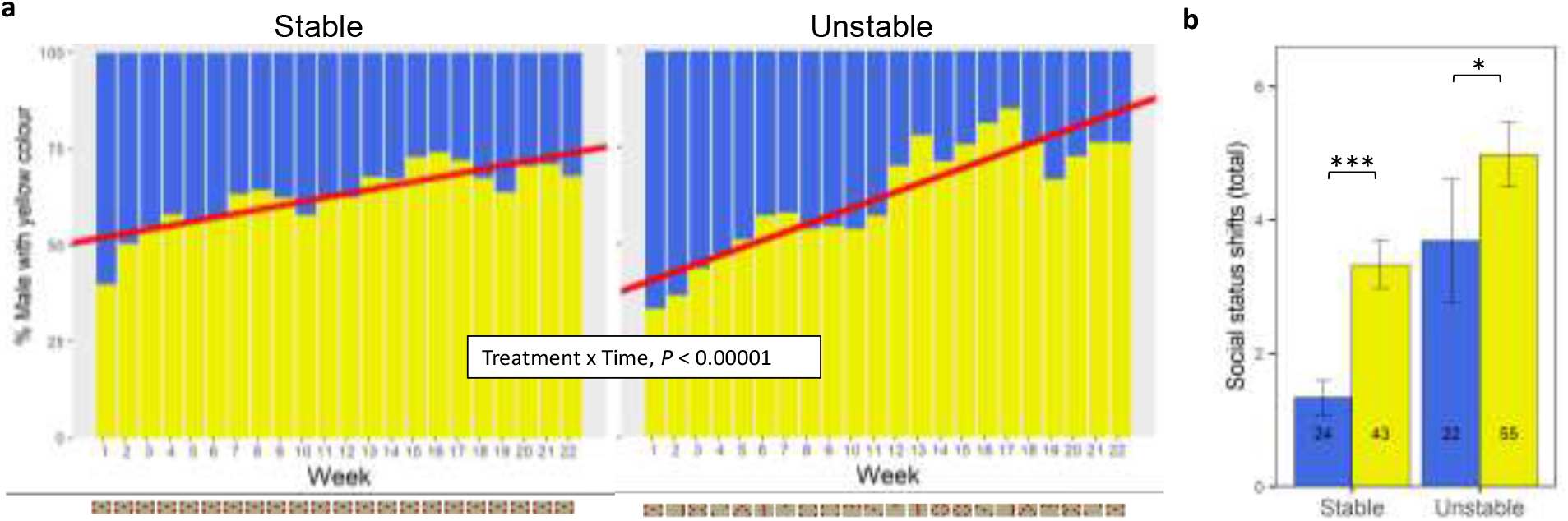
The effect of social instability on color phenotype. **a**, The percentage of yellow males over time in stable communities and unstable communities. Shown are the weekly percentages based on all communities combined (7 stable; 8 unstable communities). At the bottom is shown a schematic representation of the weekly arrangement of terracotta pot shards (top view) during the experiment. **b**, The number of social status shifts in males that were yellow or blue at the end of the experiment in stable and unstable communities. Data represented as mean ± s.e.m with sample sizes shown in the plot. Statistical findings are reported in **Extended Data Table 2**. * *P* < 0.05; *** *P* < 0.001.

Based on these findings, we hypothesized that yellow males have more success in male contest competition than blue males. Accordingly, yellow males and blue males were more likely to gain or lose territorial status, respectively, during specific time intervals (**Fig. 3a**). To formally test the advantage of yellow males in male-male competition, we staged dyadic contests between size-matched yellow males and blue males. As expected, yellow males won significantly more fights in contest competition than blue males (69.0% of contests, binomial test, *P* = 0.031; **Fig. 3b**). Given that yellow males have an advantage in male contest competition, there should be a context that favors the blue color phenotype for plasticity to persist. We therefore predicted that blue males are considered more sexually attractive than yellow males. To test this, we used binary mate choice tests and found that females spent significantly more time (percentage of time spent with blue: 74.8% ± 7.6, WSR, *P* = 0.016) and spawned more (91.7%, binomial test, *P* < 0.000001; **Fig. 3b**) with blue males than with yellow males. Collectively, our results demonstrate that alternate color phenotypes face differing trade-offs relative to male-male competition and female mate choice, which could contribute to the persistence of phenotypic plasticity in body coloration in *A. burtoni*.

**Fig. 3.**
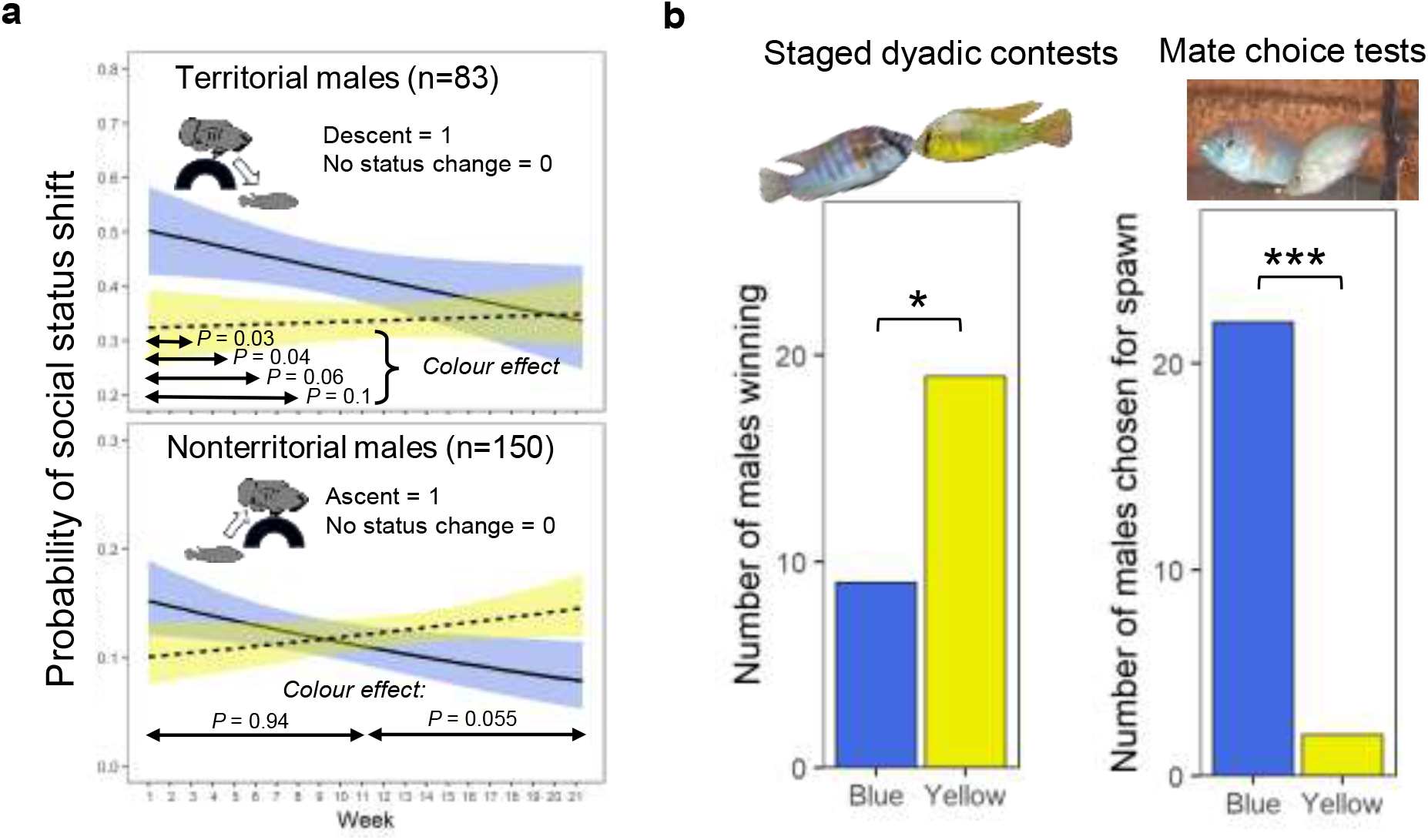
Color phenotypes and performance in male contest competition and mate choice. **a**, The probability of males changing status one week later for blue (solid line) and yellow males (dashed line) over time for territorial (top) and nonterritorial males (bottom). Specifically, the graph at the top is showing the probability of territorial males losing territorial status, with significance indicated for the color phenotype effect on this probability at varying time intervals (*P* values shown in figure indicate effect of color phenotype from separate logistic regression models for the first 2, 4, 6, or 8 weeks). The graph at the bottom shows the probability that nonterritorial males gain territorial status. Shown is the significance level of the color phenotype effect on this probability for the first and the second half of the experiment (*P* values shown in figure indicates the effect of color phenotype from logistic regression models for the first or the second half of the experiment). The analysis was restricted to males in unstable communities only. Statistical findings are reported in **Extended Data Table 3. b**, The number of blue and yellow males winning staged dyadic contests (left) and the number of blue and yellow males with whom females spawned in mate choice experiments (right). * *P* < 0.05; *** *P* < 0.001.

Oxidative stress, which is defined as an excess of reactive oxygen species relative to antioxidant capacity, can negatively impact Darwinian fitness and mediate life history trade-offs between territoriality and somatic maintenance (33, 34). To test whether the oxidative cost of male-male competition was different between yellow and blue phenotypes, we measured several markers of oxidative stress in yellow and blue males from stable communities. We found that blue males had higher oxidative stress loads than yellow males in stable communities, especially when blue males exhibited a high rate of aggressive display (**Fig. 4**) or ascended in social status (**Extended Data Fig. 3**). Our results suggest that expressing yellow or blue coloration comes with distinct costs, which likely influence the adaptive function of context-dependent color expression in *A. burtoni*.

**Fig. 4.**
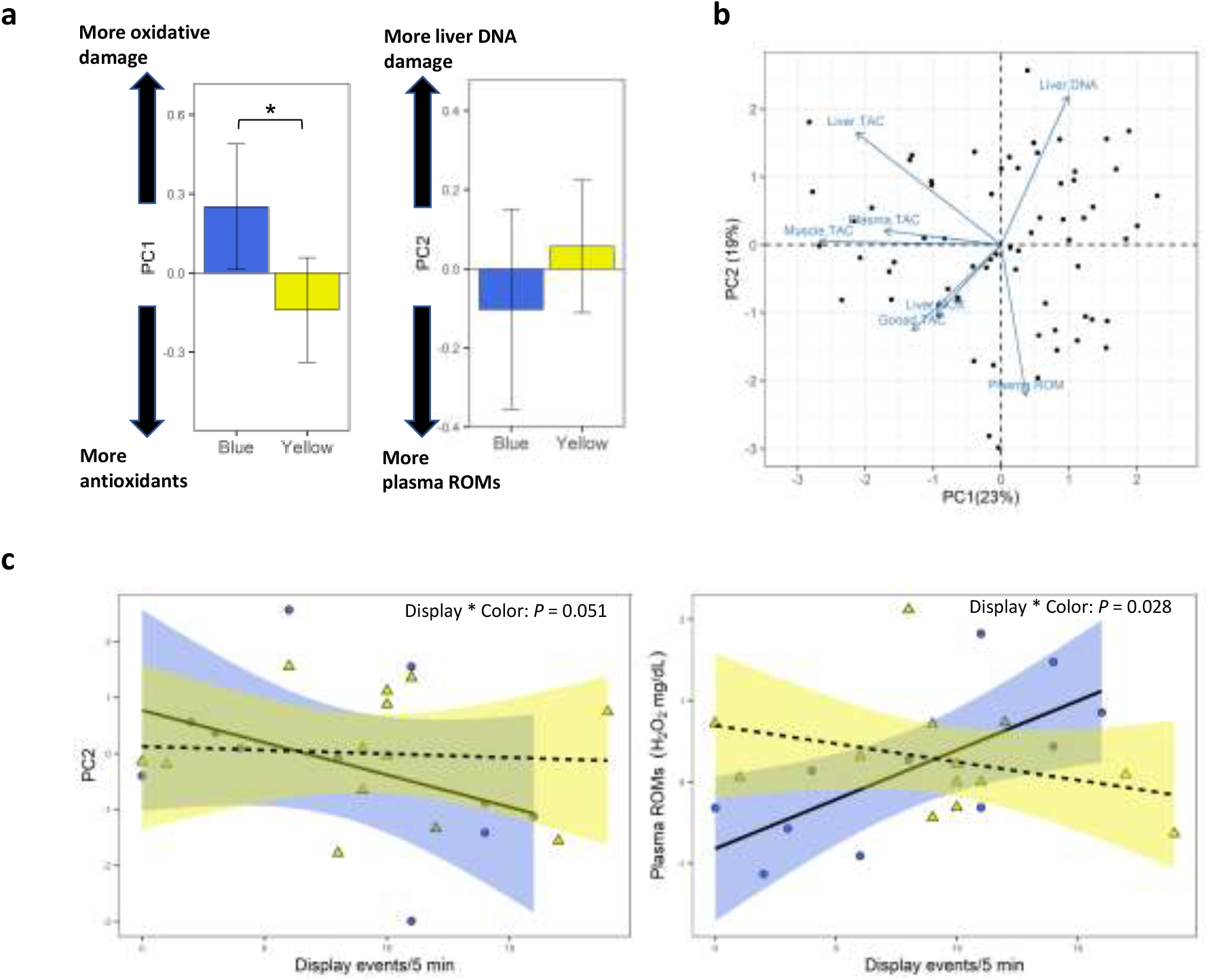
Oxidative stress profiles in blue and yellow color phenotypes. **a**, Oxidative stress profile based on the first two principal components (PC1 and PC2) for blue and yellow males. Note that higher values for PC1 correspond to higher oxidative stress (more oxidative damage and less total antioxidant capacity). This analysis was restricted to males from stable communities (24 blue males and 43 yellow males) due to the lack of blue males in unstable communities. Statistical findings are reported in the *Supplementary Information, Results*. **b**, Oxidative stress profile was based on a principal component analysis of 8 measurements of oxidative stress: total antioxidant capacity (TAC), plasma reactive oxygen metabolites (ROMs), and oxidative DNA damage. **c**, The relationship between oxidative stress profile (PC2) and territorial display rate (border display and lateral display) for blue and yellow males (left). The borderline significant interaction was mostly driven by a color phenotype dependent effect of territorial display rate on plasma reactive oxygen metabolites (ROMs), an overall measure of oxidative stress (right). Circles are blue males and triangles are yellow males. Regression lines are shown for blue males (solid lines) and yellow (dashed lines). Statistical findings are reported in the *Supplementary Information, Results*.

## Discussion

Understanding how phenotypic plasticity can persist is challenging given the costs and constraints of plasticity, such as the difficulty for animals to assess environmental cues and subsequently make timely and appropriate phenotypic adjustments (19). Factors that can promote plasticity in animal coloration have often been sought outside the realm of sexual selection, such as the ecological context favoring alternative color types (e.g. crypsis and thermotolerance (17)). Our results provide two important insights into the role of sexual selection in promoting plasticity in sexually selected traits. First, we provide experimental evidence that increased male-male competition for territories during social instability promoted the expression of yellow body coloration. We suggest that social instability creates social stress thereby altering the physiological state and coloration of the individual, possibly via the melanocortin system which is known to impact stress response, energy balance, and pigmentation (12, 35). Observed differences in oxidative stress between male color phenotypes lends some support to the idea that the physiological state is linked to body coloration, but further studies are needed to test this connection.

Second, we found that this competition-dependent color adjustment was adaptive when competing for mates. Given the importance of territoriality for reproductive success and evidence that high-quality territories can override female mate preference for nuptial coloration in cichlids (36), being a yellow morph may be adaptive in more unstable or competitive situations when being a superior fighter can aid in high-quality territory defense and acquisition. However, under stable or low-competition conditions, blue males might be able to acquire a territory while strongly benefitting from expressing the sexually attractive color phenotype.

It has been proposed that plastic polymorphisms may spark diversification when plastic traits are genetically assimilated during adaptive divergence. Recently, this hypothesis was applied to sexually selected traits since those traits are often plastic. *A. burtoni* is a riverine generalist that inhabits Lake Tanganyika and associated river systems. It is phylogenetically situated either between the Tropheini radiation and the Lake Malawi radiation (37), or between the latter and the Lake Victoria radiation (38). Given this basal position of *A. burtoni* to other species-rich cichlid clades, *A. burtoni* has been used as a model system to explore early phases of adaptive divergence by studying plastic responses in fitness-related traits (15, 39), including body coloration (12, 30, 32). The fact that yellow and blue color phenotypes as found in *A. burtoni* have been recapitulated in many haplochromine species pairs fits this scenario of initially plastic traits sparking adaptive divergence. However, our data does not support this scenario and instead suggest that sexual selection may promote the persistence of color plasticity in *A. burtoni*.

Differences in color-linked success in male-male competition and female mate choice has been previously described in the pygmy swordtail (*Xiphophorus pygmaeus*), a fish species with a fixed y-linked color polymorphism where males are either blue or gold (40). By contrast, in *A. burtoni* coloration was a highly plastic trait. Theoretical models suggest that such reversibility and conditional expression of mating type are highly vulnerable to invasion of individuals that adopt a behavioral strategy that is fixed for life due to the cost and limits of plasticity (28). However, the limits of plastic color expression are probably small because the developmental mechanisms underlying color change are robust (41), and therefore the expression of either yellow or blue color in *A. burtoni* does not necessarily limit the expression of the alternative color phenotype. The cost of plasticity requires further investigation relative to fitness decrement of plastic phenotypes to less plastic phenotypes as well as inherent trade-offs associated with alternative phenotypes (19). The high degree of reversibility that we observed in *A. burtoni* is favored when the environment varies substantially within an individual’s lifetime. Our habitat manipulation mimics natural conditions where strong daily winds or hippopotami (*Hippopotamus amphibius*) can alter territorial structures in shallow waters in the natural habitat of *A. burtoni* (27, 42). Although this needs further investigation, it seems plausible that these factors can result in the type of fine-grained, within generation variability in the competitive environment that would promote reversible phenotypic plasticity (43). Finally, we note that although our study focused on the role of sexual selection in favoring plasticity in body coloration, other factors may also contribute to adaptive color expression in *A. burtoni*, including background color adaptation (44).

In conclusion, we found evidence for adaptive plasticity driven by variation in mate competition. Males were able to alter their color phenotype in an adaptive manner to maximize their success in either male-male competition or mate attraction depending on environmental conditions. Sexual selection is thought to contribute to diversification and reproductive isolation and several studies have implicated that plasticity plays a key role in setting the stage for speciation by sexual selection (11–13, 45). However, our results suggest that sexual selection can hamper genetic divergence by contributing to the persistence of plasticity in sexual traits when alternative phenotypes are differentially favored based on the social context. Overall, our study provides novel insights into the interaction between phenotypic plasticity and sexual selection as it pertains to the evolution of diversity in sexually selected traits.

## Methods

Adult *Astatotilapia burtoni* were bred from a laboratory population originally derived from Lake Tanganyika, Africa. Fish of all experiments were housed in 28 °C aquaria with 12-h light/dark cycle containing gravel and terracotta pot shards to stimulate territoriality. Fish were fed a combination of cichlid flakes and granular food each morning and all experimental procedures were performed following feeding.

### Habitat manipulation experiment

We set up communities (n = 15) in 100-liter tanks each consisting of 10 males (n = 150) and 14 females. Communities were set up in one of two treatments: stable communities (n = 7) which each contained five terracotta pot shards in the same arrangement for 22 weeks; and unstable communities (n = 8) in which terracotta pot shard number and arrangement were altered weekly over 22 weeks (**Fig. 1a**). Male social status (territorial or nonterritorial) and color phenotype (yellow or blue) were recorded thrice weekly (**Fig. 1**) to quantify the proportion of yellow and blue males (**Fig. 2a**) as well as territorial tenure and the number of social status shifts (**Figs. 2b, 3a**).

Tanks were filmed weekly for five minutes to quantify aggressive and courtship behavior. Details of the experimental setup and protocol are described in the *Supplementary Information*.

### Biological assays

At the end of the 22-week experiment, we weighed each male, measured their standard length, and extracted tissues (blood, liver, muscle, and gonad). Gonads were weighed prior to freezing.We measured markers of 8 markers of oxidative stress and hormones (testosterone and cortisol) to assess differences in blue and yellow males. Assay information can be found in the *Supplementary Information* with a full list of oxidative stress markers found in **Extended Data Table 5**.

### Male contest experiment

We staged dyadic contests between size-matched yellow and blue males (n = 29) in 100-liter tanks. Prior to the experiment, we induced territorial status in each male by housing them individually with visual access to another male. Experimental tanks consisted of three equally sized sections. For each trial, an experimental yellow male and blue male were placed in the end compartments while a placeholder male was placed in the middle compartment. After four days in this setup, the dividers and placeholder male were removed, allowing the yellow male and blue male to engage in physical contest. Details of the experimental setup and protocol are described in the *Supplementary Information*. We recorded chases, fleeing behavior, and display behavior performed by both fish. A winner was declared upon four consecutive chases by a given fish or whichever fish was actively defending the central territory after one hour (**Fig. 3b**).

### Mate choice experiment

We measured female mate choice using a paired mate choice paradigm. Experimental tanks were identical to the male contest experiment above with the exception that the compartments were separated by dividers with holes that allowed females to move freely across sections, but the blue and yellow males were retained in the side compartments. For each trial, a gravid female was allowed to choose between a size-matched yellow male and blue male (24 females were successfully tested, each with a unique male pair). We continuously video recorded the experimental tanks from the time the female was added until the presence of eggs was detected in the female’s mouth indicating the female had spawned. Using the videos, female mate choice was determined by (i) association time (location of the female over the 24 hours prior to spawning) and (ii) final spawning preference (with whom the female physically spawned, **Fig. 3b**). For full details see *Supplementary Information*.

### Statistics

To test for the effect of social instability on color expression, we compared the proportion of yellow males in unstable and unstable communities using generalized linear mixed model (GLMM) with community as random effect (**Extended Data Table 2a**). We also tested whether blue males and yellow males at the end of the 22-week experiment differed in the number of status shifts experienced during the entire duration of the experiment (**Extended Data Table 2b**). The effect of color phenotype on behavior, hormones, and oxidative stress were performed in stable communities only since we only had enough blue males and yellow males in these communities at the end of the experiment (**Extended Data Tables 4, 5, Figs. 2, 3**). We tested which male phenotype won more fights or spawned more frequently using binomial tests (**Fig 3b**). All statistical tests were two-tailed unless stated otherwise. Additional details can be found in the *Supplementary Information*. Data and code that supports the findings of this study have been archived in Figshare: https://doi.org/10.6084/m9.figshare.21254031.v1

## Supporting information

Supplemental Methods and Results

## Acknowledgments

We thank Isaac Miller-Crews, Martinus Huigens, Jelle Boonekamp, and members of the Dijkstra lab for providing helpful comments to earlier drafts of the manuscript. Deric Learman, Francois Criscuolo, and Benjamin Swarts provided technical support. We thank Kevin Pangle, Jon Kelty, Suzy Renn, and Sebastian Alvarado for insightful discussions about this project. This research was supported by Central Michigan University, College of Science and Engineering, and the Institute for Great Lakes Research. All animal care procedures were approved by Central Michigan University Institutional Animal Care and Use Committee (IACUC protocol 15-22).

## Contributions

T.R.F., T.J.P., and P.D.D. designed the habitat disruption experiment and collected data. R.J.F. designed the mate choice experiment and collected data. S.E.B. collected data. P.M.A and R.J.F. designed the male contest experiment and collected data. P.D.D. and T.R.F. carried out the statistical analyses. P.D.D. drafted the manuscript, and R.J.F and T.R.F. contributed to writing and revision.

## Competing interests

The authors declare no competing interests.

## Notes

https://doi.org/10.6084/m9.figshare.21254031.v1

### Competing Interest Statement

The authors have declared no competing interest.

