## Supplemental Figures for "Variation in mate competition favors phenotypic plasticity in male coloration of an African cichlid"

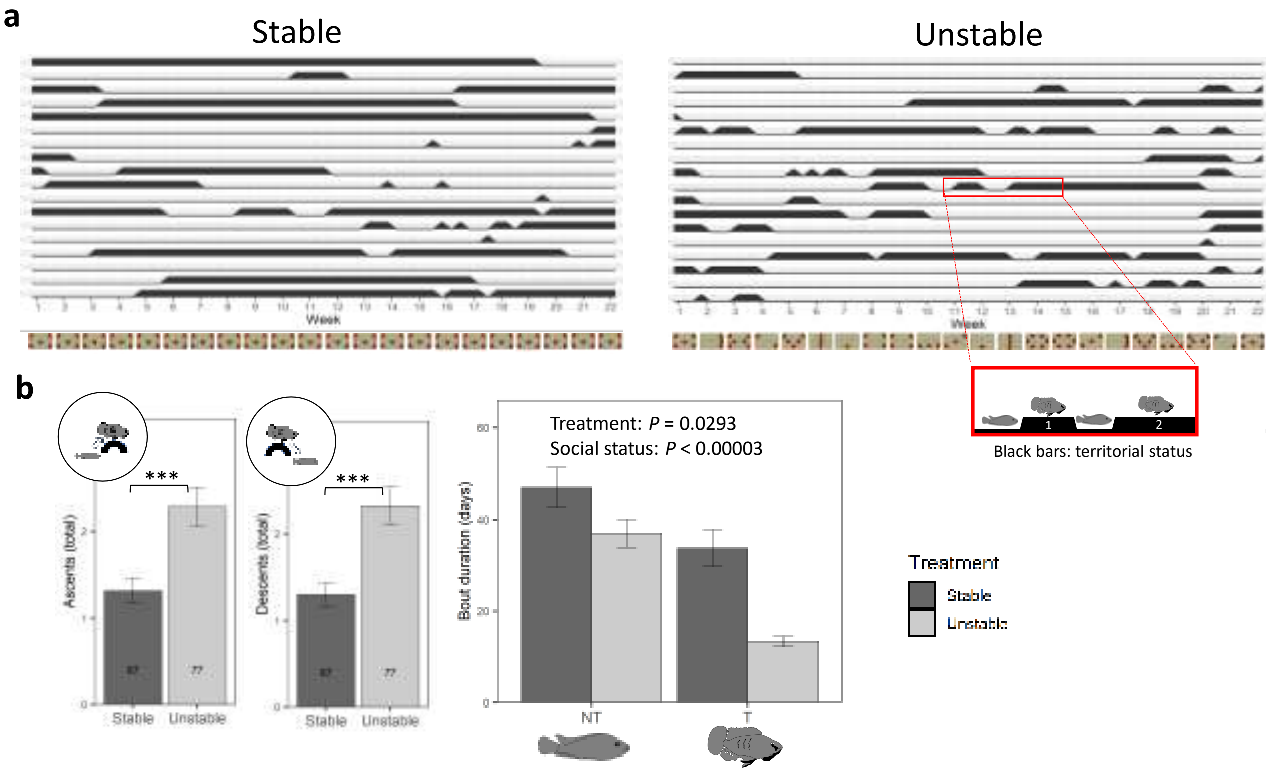


**Extended Data Fig. 1. The effect of habitat manipulation on social instability. a,** Shown is a graphical representation of social status over the entire duration of the experiment for a subset of males from stable and unstable communities. Each row represents an individual male's social status during the 22-week experiment. Black bars indicate territorial status and thin lines nonterritorial status. The timeline for the habitat manipulation is shown at the bottom with a schematic representation of the arrangement of terracotta pot shards (top view) from week 1 to week 22 in stable (n=7) and unstable communities (n=8). **b,** The frequency of social status shifts in stable communities and unstable communities separated by social ascents (gaining territoriality) and social descents (losing territoriality). Numbers in bar represent total number of males in each treatment. Data represented as mean ± s.e.m. Statistical findings are reported in **Extended Data Table 1**. *** $P < 0.001$.

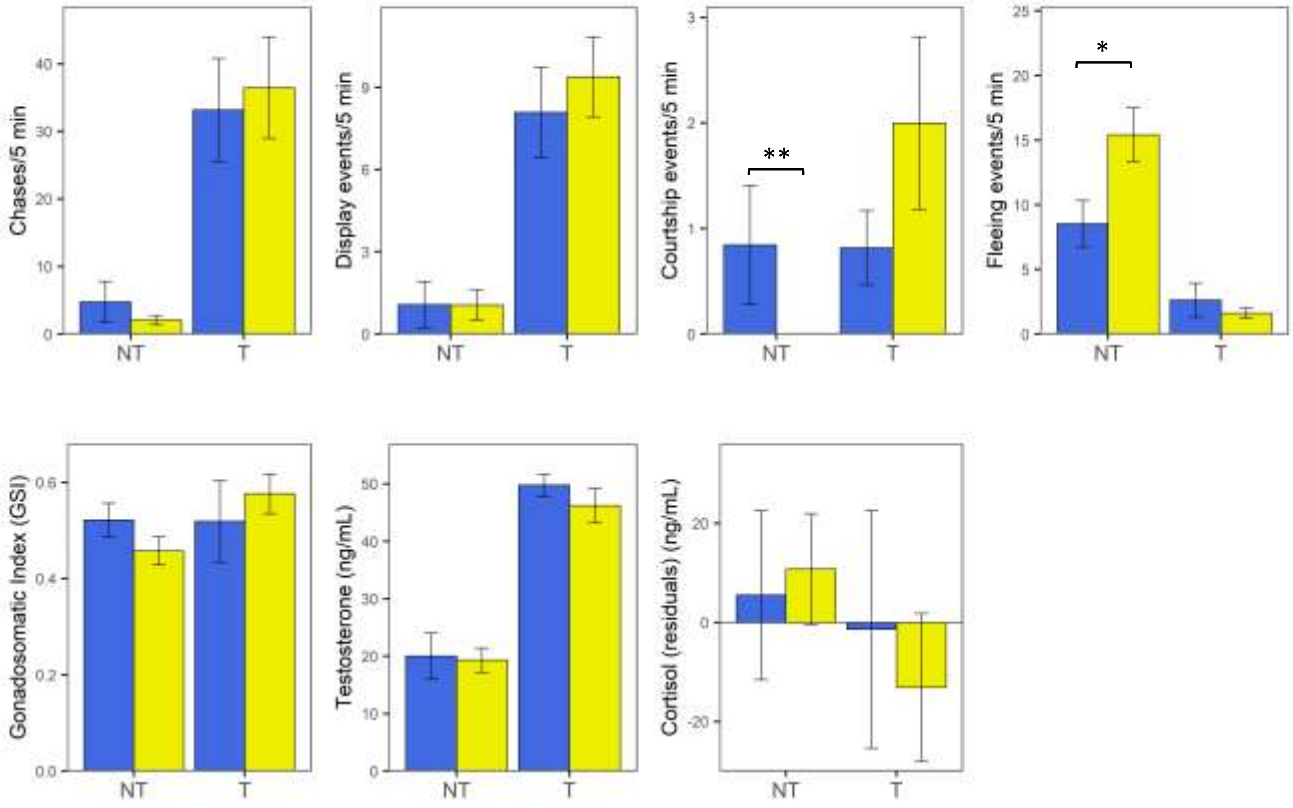


**Extended Data Fig. 2. Behavior, gonadosomatic index, and hormone levels in yellow and blue males.** Shown are the data for 24 blue and 43 yellow males from stable communities (there were not enough blue males in unstable communities for statistical comparison between color phenotypes). Data represented as mean ± s.e.m. Statistical findings and sample sizes are reported in **Extended Data Table 4**. * $P < 0.05$; ** $P < 0.01$.

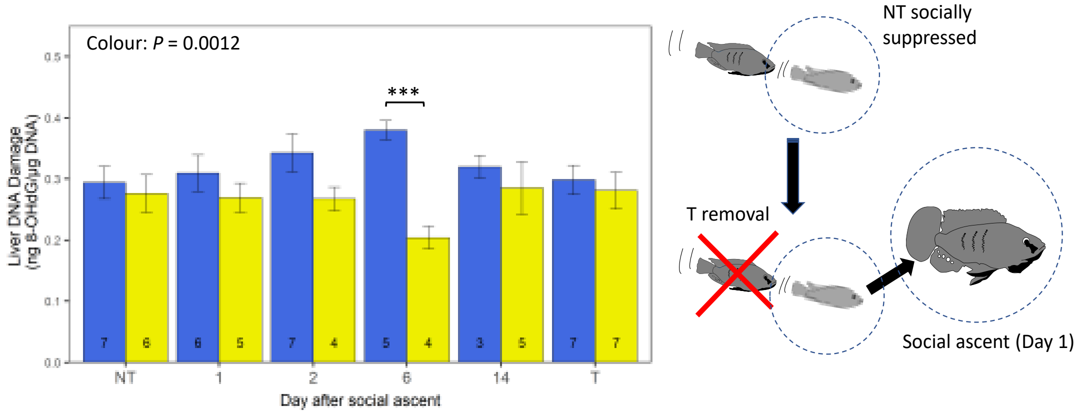


**Extended Data Fig. 3. Oxidative DNA damage during social ascent in yellow and blue males.** We used data from a separate, published dataset to examine how color phenotypes differ in oxidative stress during induced social ascent from nonterritorial (NT) to territorial status (T). Oxidative stress profile based on 7 markers of oxidative stress tended to be different between color phenotypes in males that ascended to territorial status, although this effect was not significant (day 1, 2, 6 and 14, overall color effect on PC2, LM: $0.53 \pm 0.31$, $t=1.725$, $P = 0.09$; no color effect on PC1, LM: $0.36 \pm 0.35$, $t=1.029$, $P = 0.31$; data not shown); this nonsignificant tendency was mostly driven by significantly higher oxidative DNA damage in ascending blue males compared to their yellow counterparts (overall color effect, LM: $-0.071 \pm 0.020$, $t = -3.47$, $P = 0.0012$), especially on day 6 (significance shown is based on Tukey post-hoc test). *** $P < 0.001$.
