## Supplemental Tables for "Variation in mate competition favors phenotypic plasticity in male coloration of an African cichlid"

**Extended Data Table 1**

**The effect of the habitat manipulation on social instability.** Shown are the effect of instability treatment on social status shifts (ascents and descents) and status bout duration (67 males in stable and 77 males in unstable). Shown are the model results of GLMMs. * $P < 0.05$, ** $P < 0.01$, *** $P < 0.001$

| Variable | Factor | Estimate | z value | P value |
|---|---|---|---|---|
| # Ascents | Treatment | 1.06 ± 0.35 | 3.02 | **0.0025 ** ** |
| # Descents | Treatment | 1.19 ± 0.40 | 2.97 | **0.00302 ** ** |
| # Social Status shifts | Treatment | 0.75 ± 0.21 | 3.48 | **0.0005 *** ** |
| Status bout duration[1] | Treatment | -0.18 ± 0.08 | -2.18 | **0.0293 * ** |
| | Social status | -0.30 ± 0.07 | -4.26 | **0.00003 *** ** |

[1]*This is the duration of uninterrupted bouts of territorial or nonterritorial status*

**Extended Data Table 2a**

**The effect of stability treatment on changes over time in the proportion of yellow males.** Color phenotype and social status were recorded 3 times per week during a 22-week period. Shown is the model result of a GLMM. * $P < 0.05$, *** $P < 0.001$

| Variable | Factor | Estimate | z value | P value |
|---|---|---|---|---|
| Proportion yellow | Time | 0.008 ± 0.001 | 8.77 | **< 0.00001 *** ** |
| | Treatment | -0.639 ± 0.294 | -2.17 | **0.0299 *** |
| | Time x Treatment | 0.010 ± 0.001 | 7.61 | **<0.00001 *** ** |

**Extended Data Table 2b**

**The effect of final color phenotype and stability treatment on the frequency of past social status shifts experienced during the 22-week experiment.** Final color phenotype is the color of a male at the end of the experiment. Shown is the model result of a GLMM. ** $P < 0.01$, *** $P < 0.001$

| Variable | Factor | Estimate | z value | P value |
|---|---|---|---|---|
| # Social status shifts | Final color | 0.62 ± 0.17 | 3.727 | **0.000194 *** ** |
| | Treatment | 0.36 ± 0.14 | 2.623 | **0.008714 **** |

**Extended Data Table 3**

**The probability of social status change based on color phenotype.** This analysis was restricted to unstable communities due to more opportunities for social status change. The probability of social status one week later was examined for blue and yellow males. For example, if a male was yellow in week 2, what was his probability of changing social status in week 3? Shown is the effect of color phenotype on the probability that territorial males lose territorial status (top) or nonterritorial males attain territorial status (bottom) at different time intervals. The statistical results are based on logistic regressions (GLMMs). * $P < 0.05$, ** $P < 0.01$, *** $P < 0.001$

***Probability of territorial males to lose territorial status (descent)***

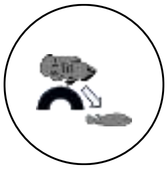

| Factor | Estimate | z value | P |
|---|---|---|---|
| Time | 0.0043 ± 0.0031 | -1.417 | 0.156 |
| Color | -0.9234 ± 0.3567 | -2.589 | **0.0096 **** |
| Time x Color | 0.0086 ± 0.0038 | 2.261 | **0.024 *** |
| *First four weeks[1]:* | | | |
| Color | -1.5392 ± 0.7533 | -2.043 | **0.041 *** |

***Probability of nonterritorial males to gain territorial status (ascent)***

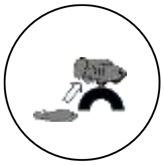

| Factor | Estimate | z value | P value |
|---|---|---|---|
| Time | -0.0043 ± 0.0026 | -1.627 | 0.104 |
| Color | -0.4405 ± 0.2577 | -1.709 | 0.087 |
| Time x Color | 0.0080 ± 0.0033 | 2.451 | **0.0143 *** |
| *First half of the experiment:* | | | |
| Color | -0.0180 ± 0.2300 | -0.078 | 0.94 |
| *Second half of the experiment:* | | | |
| Color | 0.6108 ± 0.3189 | 1.916 | 0.055 |
| Time | 0.0150 ± 0.0043 | 3.457 | **0.0005 ***** |

[1] *The analysis yielded similar results for the first 2 or 6 weeks, see Fig. 3 for P values.*

**Extended Data Table 4**

**Behaviour, gonadosomatic index (GSI), and hormone levels in yellow and blue males.** This analysis was restricted to stable communities only because there were not enough blue males in unstable communities at the end of the experiment. We used ‘final color’ (coloration at the end of the 22-week experiment) as an independent variable and tested it in combination with social status (territorial (T) or nonterritorial males (NT)). If model assumptions were violated, we implemented separate tests within each category of male (T or NT). Shown is the variable tested, sample sizes for blue and yellow males, and category of males used in each statistical test. Shown are the statistical results of LMMs (with t statistic), GLMMs (with z statistic) or nonparametric Mann-Whitney U tests (with W statistic). * $P < 0.05$, ** $P < 0.01$

| **Variable** | **#Blue, #Yellow males**[1] | **Category** | **Estimate** | **Statistic** | **P value** |
|---|---|---|---|---|---|
| Chases | 13, 30 | NT | | W = 196 | 0.99 |
| | 11, 13 | T | 0.11 ± 0.31 | z = 0.363 | 0.72 |
| Display events | 13, 30 | NT | | W = 189.5 | 0.86 |
| | 11, 13 | T | 0.09 ± 0.28 | z = 0.319 | 0.75 |
| Courtship | 13, 30 | NT | | W = 240 | **0.008 **** |
| | 11, 13 | T | -0.028 ± 0.822 | z = -0.034 | 0.973 |
| Fleeing | 13, 30 | NT | 0.619 ± 0.249 | z = 2.489 | **0.0128 *** |
| | 11, 13 | T | | W = 70.5 | 0.976 |
| GSI | 22, 43 | Combined[2] | -0.022 ± 0.046 | $t_{65.67} = -0.486$ | 0.629 |
| Testosterone | 24, 42 | Combined[3] | -0.066 ± 0.084 | z = -0.79 | 0.432 |
| Cortisol | 24, 41 | Combined[2] | 1.409 ± 15.78 | $t_{65} = 0.089$ | 0.929 |

[1]*Sample sizes may vary depending on availability of sample.*
[2]*Social status was not included in this model because it was not significant.*
[3]*Social status was included in this model as a significant effect.*

**Extended Data Table 5**

**Statistical comparisons of measurements of oxidative stress between yellow and blue males.** This analysis was restricted to stable communities only because there were not enough blue males in unstable communities at the end of the experiment. Social status did not influence the results (Ps > 0.2). Shown are the sample sizes and the model results of LMMs (with t statistic) or GLMMs (with z statistic) using 'final color' (coloration at the end of the 22-week experiment) as independent variable.

| Variable | #Blue, #Yellow males[1] | Estimate | Statistic | P value |
|---|---|---|---|---|
| Plasma ROMs | 24, 42 | -0.43 ± 0.50 | $t_{66} = -0.859$ | 0.394 |
| Plasma TAC | 24, 41 | 223.26 ± 144.41 | $t_{63.9} = 1.546$ | 0.127 |
| Gonad TAC | 23, 43 | 0.081 ± 0.049 | $t_{65.98} = 1.641$ | 0.106 |
| Muscle TAC[2] | 24, 43 | 0.017 ± 0.042 | $t_{62.63} = 0.412$ | 0.682 |
| Liver TAC | 24, 43 | 0.024 ± 0.030 | $t_{60.79} = 0.786$ | 0.434 |
| Liver DNA damage | 21, 38 | -0.040 ± 0.054 | z = -0.750 | 0.453 |
| Liver NOX | 18, 37 | -0.115 ± 0.211 | z = -0.548 | 0.584 |

[1]*Sample sizes may vary depending on availability of sample.*
[2]*This model also included the effect of social status (-0.09 ± 0.05, P = 0.058) and weight (0.026 ± 0.009, P = 0.004).*
*ROMs: reactive oxygen metabolites; TAC: total antioxidant capacity; DNA damage: oxidative DNA damage based on 8-OHdG; NOX: NADPH-oxidase activity (this enzyme can generate reactive oxygen species).*
