## Supplemental Methods and Results for "Variation in mate competition favors phenotypic plasticity in male coloration of an African cichlid"

**METHODS………………………………………………………………………………2**
**RESULTS……………………………………………………………………………….10**

### METHODS

#### STUDY ANIMALS AND EXPERIMENTAL PROTOCOL

For this experiment adult *Astatotilapia burtoni* were bred from a laboratory population originally derived from Lake Tanganyika, Africa (1). Fish were housed in aquaria kept at 28 °C with a 12-h light/dark cycle and 10 minute each dusk and dawn period to mimic natural settings. Aquaria contained gravel substrate and terracotta pot shards to stimulate territoriality. Fish were fed a combination of cichlid flakes (Omega Sea LLC, Painesville, Ohio) and granular food (Allied Aqua, Smithville, Missouri) each morning between 8 and 10 am. All experimental procedures and observations were performed in the morning (unless indicated otherwise) at least 10 minutes after feeding. Continuous water flow and central mechanical and biological filtration occurred throughout the entirety of the experiment. Experimental fish were individually tagged through the dorsal musculature using a stainless-steel tagging gun and colored beads. All animal care procedures were approved by Central Michigan University Institutional Animal Care and Use Committee (IACUC protocol 15-22) and were in compliance with the US National Research Council's Guide for the Care and Use of Laboratory Animals, the US Public Health Service's Policy on Humane Care and Use of Laboratory Animals, and Guide for the Care and Use of Laboratory Animals.

#### HABITAT MANIPULATION EXPERIMENT

To examine how social instability affects the expression of blue and yellow coloration, we set up communities (n = 15) in 100-liter tanks (76 x 51 x 30 cm) each consisting of 10 males and 14 females (total of 150 males, mass = 7.02 ± 0.15 g, standard length = 61.9 ± 0.47 mm). Each community had one terracotta pot shard placed in each corner of the tank and one terracotta pot shard placed in the center of the tank. After their formation, communities were given 4 weeks to settle before the experiment began. Territorial males typically defend one terracotta pot shard and removing, adding, or rearranging the terracotta pot shards can lead to loss of territoriality or a new male gaining territoriality. Treatments consisted of 7 stable communities with unchanging territory arrangements and 8 unstable communities in which terracotta pot shard number and arrangement were altered weekly as shown in **Fig. 1**. The first manipulation took place at the end of week 1 and the experiment lasted a total of 22 weeks. In stable communities, terracotta pot shards were removed and then immediately replaced in their previous location to account for handling stress. During the 22-week experiment, a total of 6 out of 150 males died (mortality rate of 4%), which did not appear to be driven by treatment, social status or color phenotype. Males that died were replaced, except for one male which died just prior to tissue collection. Replacement males were excluded from the analysis.

*Scoring of social status and color phenotype*: To quantify shifts and tenure of male color and social status, we recorded social status and color phenotype for each male three times per week (Monday, Wednesday, and Friday), with the first observation starting at the beginning of week 1.

The third observation in each week occurred just prior to the habitat manipulation (that is, three observations were done before the first habitat manipulation). During each observation, we quantified color phenotype as being yellow or blue as previously described (2). Social status was assigned by characterizing males as territorial (dominant) or nonterritorial (subordinate). Territorial males were defined as guarding a terracotta pot and engaging in regular aggressive behaviors with other males (chasing, border displays, and lateral displays) as well as chasing and courting females. Nonterritorial males were characterized as not defending a territory, schooling with females, and rarely engaging in aggressive behaviors with other males (3).

*Behavior*: To allow for behavioral analysis, tanks were video recorded weekly for five minutes just prior to the habitat manipulation. We quantified the behavior of all fish (n = 144) using the last five videos taken before tissue collection. Behavior was quantified using focal observations. We recorded the number of chasing and display behaviors (border displays and lateral displays) as well as the number of fleeing events. We also recorded courtship behavior, which typically consisted of a lead swim combined with a quiver. Behavioral definitions have been described previously (4).

*Oxidative stress*: At the end of the 22-week period, males were weighed and their standard length was measured before being euthanized and collecting their tissue as described previously (4). Gonads were weighed prior to freezing. Oxidative stress, which is defined as an excess of reactive oxygen species relative to antioxidant capacity, can negatively impact Darwinian fitness and mediate life history trade-offs between territoriality and somatic maintenance(5). Therefore, we compared several markers of oxidative damage and antioxidant capacity in blue and yellow males. We measured plasma reactive oxygen metabolites (ROMs) as a measure of overall oxidative damage. Plasma total antioxidant capacity (TAC) was measured as an indicator of overall antioxidant protection. Oxidative DNA damage (based on 8-hydroxy-2′-deoxyguanosine (8-OhDG) content) was measured as a more specific marker of oxidative damage. Nicotinamide adenine dinucleotide phosphate (NOX) activity was assessed as an indirect marker of reactive oxygen species. DNA damage and NOX activity were measured in the liver because metabolic processes in the liver often vary with life history stage. In addition to plasma, TAC was also measured in liver, muscle, and gonad tissue to evaluate antioxidant capacity in important tissue types. For detailed protocols, see Border et al. 2021 (6).

*Hormones*: To compare hormone levels between yellow and blue males we measured circulating cortisol and testosterone levels using ELISA kits (Enzo Life Sciences) as previously described (6). The intra-assay CV were 2.9% and 3.3% for cortisol and testosterone, respectively. The inter-assay CV were 2.7% and 2.9% for cortisol and testosterone, respectively.

**MALE CONTEST EXPERIMENT**

To test which color phenotype has an advantage in direct male contest competition, we staged dyadic contests between yellow and blue males as described below (modified from (7)).

*Pre-experimentation housing*: Males were housed in four equally sized individual compartments using clear perforated dividers placed in 100-liter tanks (76 x 51 x 30 cm) for 1-2 months prior to experimentation exactly as described elsewhere (2) (n = 58, mass = 14.40 ± 0.33 g, standard length = 79.99 ± 0.70 mm). Males had visual access to at least one neighboring male to stimulate territoriality and to avoid unwanted effects of social isolation while preventing physical interaction (8). Standard length and weight of males were measured at least a week prior to use in experimentation to form blue-yellow pairs. We made sure pairs were mostly size matched (maximum size difference was less than 13 mm) such that size asymmetry was balanced with respect to color phenotype. To avoid bias, isolated neighbors were never chosen as experimental pairs for tests of dominance.

*Staged combat*: Male pairs were placed in an experimental 100-liter tank (76 x 51 x 30 cm) consisting of three equally sized sections separated with clear horizontal dividers. A terracotta pot shard was placed in each compartment to serve as the focal point for territorial defense. The middle portion of the tank housed a placeholder male, while the two compartments on either side housed an experimental male, each of which had opposing colors (i.e., if a male on one side was yellow, the male in the opposing side was blue). The placeholder male facilitated the acclimation of experimental males to the experimental tank. Males were allowed four days to acclimate to the experimental tank. Between 4:00 pm and 5:00 pm on the fourth day of acclimation to the experimental tank, the placeholder male was removed from the center compartment. The following morning, males were fed and after at least ten minutes the dividers and the terracotta pot shards in the experimental male compartments were removed simultaneously. The experimental males were then allowed to engage in full-contact physical combat over ownership of the remaining central terracotta pot shard. We recorded chases, fleeing behavior, and display behavior performed by both fish. Winners of each combat trial were declared when one fish chased the other four consecutive times. If no winner was declared in fifteen minutes, the tank was observed after one hour to discern which fish was the winner, defined as the male who was actively chasing the other fish and occupying the terracotta pot shard. Each experimental fish had its weight and standard length measured immediately after the combat trial. Males within pairs did not differ in weight (paired t-test, $t_{27} = -0.425$, $P = 0.67$) or standard length ($t_{27} = -1.255$, $P = 0.22$). We staged fights between 29 male pairs and eliminated one fight in which a male changed color before onset of the fight. All fights resulted in a winner and a loser as defined above.

### MATE CHOICE EXPERIMENT

To test which color phenotype is preferred by females, we used a paired mate choice paradigm based on behavioral preference (time spent with) and actual spawning behavior.

*Pre-experimentation housing*: Prior to use in mate choice trails, males (referred to as 'stimulus males') were housed in individual compartments for at least 2 weeks as described for the pre-experimentation housing in the male contest experiment (n = 64, mass = 14.95 ± 0.52 g, standard length = 80.00 ± 0.85 mm). The pre-experimentation housing condition ensured territorial status and full expression of body coloration in each stimulus male. Experimental females were housed in female-only 100-liter tanks (76 x 51 x 30 cm) consisting of 15-25 females. Females were selected from female-only holding tanks for experimental trial when gravid as characterized by distended abdomen (9) (n = 32, mass = 4.60 ± 0.25 g, standard length = 54.57 ± 1.00 mm).

*Mate choice setup.* The 100-liter experimental tanks were separated into three equal horizontal sections each containing a terracotta pot shard. The clear dividers were perforated with 2-centimeter holes that are large enough for females to swim freely across compartments but too small for males to pass through, similar to a protocol described previously (10). The middle portion of the tank housed a placeholder male, while the two compartments on either side each housed a stimulus male. Stimulus males were allowed to settle in for 7-10 days before replacing the placeholder male with a gravid female.

*Procedure.* In each mate choice trial, a female was given a choice between a unique, size matched blue and yellow stimulus male. The size difference between stimulus males in each pair was less than 5 mm and we ensured that the size asymmetry with respect to male color phenotype was balanced. In each trial, the experimental female was added immediately after the placeholder male had been removed (within five minutes). The behavior of all fish was video recorded continuously using a 1080p Yi camera. Recording occurred until the female began holding eggs in her mouth. We checked whether the female had spawned by observing the experimental female 1-2 times per day and stopped the video recording when the female was mouthbrooding. After spawning, the female was netted and the eggs were removed and counted. We started 32 mate choice trials and successfully tested mate preferences in 24 females (57.04 ± 4.77 eggs per spawning). Two females did not spawn, and six trials were aborted because one or both stimulus males changed color phenotype. Most females spawned within 5 days after being released into the mate choice compartment (3.7 ± 0.6 days). Each experimental fish had its weight and length measured immediately after the spawning was recorded. Stimulus males within pairs did not differ in weight (paired t-test, $t_{23} = 0.48132$, $P = 0.63$) or standard length ($t_{23} = 0.81879$, $P = 0.42$).

*Behavioral observations and analysis.* Using the videos, we determined mate preference based on spawning behavior and association time prior to spawning. Immediately before spawning, the chosen male actively displayed courtship behavior. Spawning events were characterized as a prolonged (between 10 and 30 seconds) visit to a male's terracotta pot shard by a female after an active courtship display. Spawning always took place in a dug-out spawning area near the terracotta pot shard where the male and female showed the characteristic circling behavior associated with spawning (11). There was only one female who visited both stimulus males during a spawning but upon closer examination still only laid eggs with one male. After spawning occurred, the female moved to the center compartment (duration of spawning, 394.8 ± 69.3 sec). In all cases, females were observed not carrying eggs before the spawn, and carrying eggs in their mouth after the spawn. The females also had a more concave belly after the spawn. In addition, females showed the characteristic upward lift of the anterior portion of the body after the spawn (12). Based on these observations, we were able to assign unambiguously with which stimulus male the females spawned.

To assess association time as a measure of mate preference, we recorded whether the female was in the center compartment, the left stimulus male's compartment, or the right stimulus male's compartment. The location of the female was recorded 24 hours before the spawn until the onset of spawning at five-minute intervals. We excluded the time when the lights were turned off. The experimenter recording the location of the female was blind with respect to the spawning decision of each focal female.

### STATISTICAL ANALYSIS

All analyses were conducted in R v3.6.1 (2019) (R Core Team, 2012). We analyzed our data using the R package lme4 (13) and glmmTMB (14). We used linear mixed models (LMMs, with tank as a random effect) with a maximum-likelihood protocol (R package lme4) to model our data. For count and proportional data, we used generalized linear mixed models (GLMMs) with tank as a random effect. GLMM hurdle models were also considered for count variables that contained zero values. Appropriate distributions for GLMMs were chosen based on AIC values and model diagnostics. To evaluate the validity of our LMM and GLMM models, we examined the residuals, qqplots, and plots of predicted values versus residuals. We report mean ± SE for our model estimates. All statistical tests were two-tailed unless stated otherwise.

*Effect of habitat manipulation on social stability:* When a male changed social status between two consecutive observations, it was considered an ascent (nonterritorial to territorial transition) or descent (territorial to nonterritorial transition). The sum of ascents and descents of a male was considered the number of status shifts that a male experienced. To validate that our habitat manipulation induced social instability, we tested whether changing the number and arrangement of terracotta pot shards led to more ascents and descents using a GLMM hurdle model assuming a negative binomial distribution. To test whether the habitat manipulation led to shorter status tenure (i.e. uninterrupted bouts of nonterritorial and territorial status), we compared status tenure using GLMM assuming a negative binomial distribution. Statistical results are reported in **Extended Data Table 1**, and a summarized version of significance testing is shown in **Extended Data Fig. 1**.

*Frequency of color phenotype change.* To test whether social instability increases the number of yellow males, we calculated the proportion of yellow and blue males during each global observation in each community. We then tested whether the change in proportion of yellow males over time differed between stable and unstable communities using GLMM assuming a beta distribution. Next, we tested whether the number of status shifts (sum of ascents and descents) a male experienced during the 22-week experiment predicted final color using a GLMM assuming a negative binomial distribution.

We quantified the number of color phenotype changes of each male (n = 144) during the 22-week experiment (blue to yellow or yellow to blue). To test whether the number of color phenotype changes differed between stable and unstable communities, we used a GLMM assuming a negative binomial distribution.

*Probability of social status change based on color phenotype:* Based on the observed increase in yellow males in unstable communities, we hypothesized that yellow males have an advantage in competition for territories. As an initial test of this hypothesis, we evaluated whether color phenotype predicted social status change. This was done in unstable communities only since we manipulated the habitat in these communities to trigger social status changes. To this end, we recorded for each observation whether a male (that was either yellow or blue at that moment) underwent a social status change within the next week. If a social status change occurred, it was coded as '1', and if no social status change occurred, it was coded as '0'. This was done for territorial and nonterritorial males separately. We recorded the probability of social status change using a one-week interval rather than the subsequent observation (2-3 days later) because the habitat manipulation took place weekly, and we wanted to ensure there was at least one manipulation between the recording of color phenotype and probability of social status change. To test whether color phenotype predicted social status change, we first fitted a logistic regression (GLMM assuming a binomial distribution) with time and color phenotype as interaction effect. Since the interaction effect was significant for both territorial and nonterritorial males (see main text), we tested for the effect of color phenotype on the probability of social status change focusing on relevant time intervals.

*Color effect on behavior and hormones in stable communities:* We restricted the color phenotype analysis to stable communities only because there were not enough blue males in unstable communities at the end of the experiment (one territorial blue male was present in all unstable communities combined). In stable communities there were a total of 24 blue males (11 territorial and 13 nonterritorial males) and 43 yellow males (13 territorial and 30 nonterritorial males) at the end of the experiment. We tested whether blue or yellow males differed in behavior (chases, display events, courtships, and fleeing), gonadosomatic index, and hormone levels using the following approach. For the behavior we used separate tests (LMMs, GLMMs assuming negative binomial distribution) for T and NT because the distribution of the data was very different for these two categories of males. Mann-Whitney U tests were used if assumptions for mixed models were not met.

Testosterone was analyzed using both social status and color phenotype as fixed effects using a GLMM assuming a gaussian log link function. Gonadosomatic index and cortisol (residuals to control for the effect of time of blood sampling) were analyzed using LMMs with only color phenotype as a fixed effect because social status did not improve the fit of these models. Gonadosomatic index was calculated as (gonad weight/total body weight)*100.

*Color effect on oxidative stress in stable communities:* To test whether blue and yellow males within stable communities differ in oxidative stress, we used principal component analysis (PCA) which summarizes the seven measures of oxidative stress by fewer axes. Missing values (e.g. due to insufficient tissue sample; 23 out of 446 (5.2%) of data was missing) were filled using iterative PCA algorithm using the miss.MDA package(15). Before entering the PCA, we standardized all measurements by calculating z-scores, which were obtained by subtracting the entire sample mean from each observation and dividing by the standard deviation. The first three principal components (PCs) accounted for 57% of the variation in the original dataset. We restricted further analyses to PC1 and PC2 since they captured most of the variation (eigenvalues of 1.6 and 1.3, respectively while the other principal components had eigenvalues of 1.1 or less), with PC1 explaining 23% and PC2 explaining 19% of the total variance. We then compared principal component 1 and 2 between blue and yellow males using LMMs with final color and social status as independent variables. To examine each marker independently, we also used LMMs with final color and social status as independent variables.

To test whether blue and yellow males differ in the trade-off between aggressive behavior and oxidative stress, we tested whether the relationship between aggressive behavior (chases or displays) and oxidative stress was dependent on color phenotype within territorial males in stable communities. This was done by fitting LMMs with principal component 1 and 2 as dependent variables and chases or displays with final color as interaction effects.

*Color effect on oxidative stress in ascending males:* We found that there was a difference in oxidative stress levels between blue and yellow males within stable communities (see results section). To investigate the robustness of this findings, we re-analyzed a previously published dataset where nonterritorial males were allowed to become territorial by removing the territorial male (4). This resulted in the nonterritorial male assuming territorial (dominance) status within minutes after light onset after the territorial male was removed (the territorial male was removed ~ one hour before lights came on). Males were sampled the day of social ascent (day 1) as well as 2, 6 and 14 days after social ascent. In this study, we measured Reactive Oxygen Metabolites (ROMs), blood and liver DNA damage (8-OhDG content), and liver NOX activity. Antioxidants were measured as total antioxidant capacity (TAC) in plasma and liver as well as the enzymatic superoxide dismutase (SOD) in liver tissue.

We carried out the same PCA analysis as described above and found that PC1 explained 28% and PC2 explained 20% of the total variance in the original dataset. We then compared principal component 1 and 2 between blue and yellow males using LMMs with final color and day of ascent as independent variables. To examine each marker independently, we also fit linear models with final color and day of ascent as independent variables. We used Tukey's multiple comparison post-hoc tests to compare color phenotypes within days using the emmeans package (16).

*Male contest experiments*: Size asymmetry within each pair did not influence contest outcome. We therefore tested whether blue or yellow males had an advantage in male contest using a binomial test. A one-tailed test was used because previous studies have indicated that yellow males have an advantage in dyadic contests (17) and our social status change probability analysis also supported this competitive advantage of yellow males.

*Mate choice experiments*: As noted above, males within each stimulus pair were sized matched and it was therefore not necessary to control for size asymmetry in the analysis. We first tested whether females spent more time with blue or yellow males. This was done using a Wilcoxon signed-rank test by comparing the percentage of time spent with blue against the no preference line. We then compared the number of females that spawned with blue or yellow males using a binomial test.

### RESULTS

**Figure 4, complete statistical results.** Blue and yellow males have distinct oxidative stress profiles based on principal component 1 (LMM, effect of final color: -0.529 ± 0.225, $t_{61.16} = -2.35$, $P = 0.022$) but not principal component 2 (0.006 ± 0.0.248, $t_{61.90} = 0.022$, $P = 0.982$) (**Fig. 4a**). Since higher values for PC1 reflect a combination of more oxidative damage (liver DNA damage and plasma reactive oxygen metabolites (ROMs)) and less total antioxidant capacity, blue males experienced more oxidative stress than yellow males. However, note that there were no significant differences for each measurement individually (*P*s > 0.106, **Extended Data Table 5**). This analysis was restricted to males from stable communities (24 blue males and 43 yellow males) due to the lack of blue males in unstable communities at the time of tissue collection.

Oxidative stress profile (PC2) was related to territorial display rate (border display and lateral display) in a color phenotype dependent manner (**Fig. 4b**, left), although this effect did not reach statistical significance (LMM, display * final color: 0.179 ± 0.087, $t_{22.65} = 2.06$, $P = 0.0510$) after controlling for the effect of body mass (0.333 ± 0.142, $t_{22.88} = 2.355$, $P = 0.028$). This interaction effect in PC2 was mostly driven by a color phenotype dependent relationship between territorial display rate and plasma reactive oxygen metabolites (ROMs), an overall measure of oxidative damage (**Fig. 4b**, right) (LMM, display * final color: -0.327 ± 0.102, $t_{23} = -3.194$, $P = 0.004$). Oxidative stress was not influenced by chase rate and final color (*P*s > 0.2, data not shown). Circles are blue males and triangles are yellow males. Regression lines are shown for blue males (solid lines) and yellow (dashed lines).
